# Syotti: Scalable Bait Design for DNA Enrichment

**DOI:** 10.1101/2021.11.05.467426

**Authors:** Jarno Alanko, Ilya Slizovskiy, Daniel Lokshtanov, Travis Gagie, Noelle Noyes, Christina Boucher

## Abstract

Bait-enriched sequencing is a relatively new sequencing protocol that is becoming increasingly ubiquitous as it has been shown to successfully amplify regions of interest in metagenomic samples. In this method, a set of synthetic probes (“baits”) are designed, manufactured, and applied to fragmented metagenomic DNA. The probes bind to the fragmented DNA and any unbound DNA is rinsed away, leaving the bound fragments to be amplified for sequencing. This effectively enriches the DNA for which the probes were designed. Most recently, Metsky et al. (Nature Biotech 2019) demonstrated that bait-enrichment is capable of detecting a large number of human viral pathogens within metagenomic samples. In this work, we formalize the problem of designing baits by defining the Minimum Bait Cover problem, which aims to find the smallest possible set of bait sequences that cover every position of a set of reference sequences under an approximate matching model. We show that the problem is NP-hard, and that it remains NP-hard under very restrictive assumptions. This indicates that no polynomial-time exact algorithm exists for the problem, and that the problem is intractable even for small and deceptively simple inputs. In light of this, we design an efficient heuristic that takes advantage of succinct data structures. We refer to our method as syotti. The running time of syotti shows linear scaling in practice, running at least an order of magnitude faster than state-of-the-art methods, including the recent method of Metsky et al. At the same time, our method produces bait sets that are smaller than the ones produced by the competing methods, while also leaving fewer positions uncovered. Lastly, we show that syotti requires only 25 minutes to design baits for a dataset comprised of 3 billion nucleotides from 1000 related bacterial substrains, whereas the method of Metsky et al. shows clearly super-linear running time and fails to process even a subset of 8% of the data in 24 hours. Our implementation is publicly available at https://github.com/jnalanko/syotti.

## 1 Introduction

Our understanding of microbial species has evolved at an impressive rate, but it is estimated that 99% of microoganisms cannot live outside their natural environments, and thus, cannot be cultured and sequenced [15]. This created a significant roadblock in studying such species. In the mid-2000s, metagenomic shotgun sequencing became widely available, which enabled the sequencing of all of the DNA within non-cultured samples—whether a swab of a person’s mouth or a soil sample. This enabled the large-scale study of microoganisms, and thus greatly expanded our knowledge in this field. However, many scientific questions focus on specific sequences such as antimicrobial resistance (AMR) genes [14], or human viral strains and substrains [4]. To address these questions, most of the sequence reads in a metagenomic dataset are irrelevant, and are typically eliminated from further consideration. Moreover, the percentage of such reads can be quite high; for instance in agricultural samples, 90 to 95% of reads are often eliminated because they are not of interest to the research question [14, 17, 9]. Furthermore, using shotgun metagenomics in these scenarios not only leads to undue monetary expense but also greatly lowers the sensitivity of detection. For example, Lee et al. [10] demonstrated that out of 31 viral strains identified, 11 were unidentified via traditional sequencing, and Noyes et al. [14] reported that 1,441 AMR genes within agricultural samples were undetected by standard metagenomic sequencing.

One way to address these issues is to move beyond sequencing the entire microbial population within a sample, and rather to select the sequences of interest for targeted sequencing. This can be accomplished through *targeted enrichment*, which uses *biotinylated cDNA bait molecules* to target specific regions from DNA libraries for sequencing. More specifically, baits (short synthetically created single-stranded cDNA molecules) bind to their DNA targets and are then captured within the sample using a magnet. Non-captured, unbound DNA fragments (i.e., non-targeted DNA) are then rinsed away. Thus, only bound—i.e., targeted—DNA is sequenced. Although non-targeted DNA is not completely eliminated from sequencing, it is greatly reduced. One of the first, and arguably the most critical steps within this process requires solving a computational problem: for a given set of target DNA sequences (e.g., set of genes or viral strains) and a specified bait length *k*, identify a set of baits such that there exists at least one bait in that set that binds to each position of every sequence in the database.

Initially one could expect that baits could be computationally designed by finding all unique *k*-length subsequences (*k*-mers) in the targeted DNA. This solution however is not feasible as the number of *k*-mers grows rapidly with increasing dataset size. Thus, two key challenges need to be addressed for effective bait design. First, the number of baits must be minimized, as the cost of targeted enrichment is proportional to the size of the bait set and there is an upper limit set by the bait manufacturer. For example, Agilent Technologies, Inc. sets a limit of approximately 220,000 baits; larger bait sets can be manufactured, but would need to be split into multiple sets, thus greatly increasing the cost and labor of purchasing and processing multiple bait sets per sample. Second, the baits do not bind exactly to a single DNA sequence identical to that bait, but will bind to any subsequence with some allowed number of mismatches between the bait and target DNA sequence (typically, over 70% of positions must match). For this reason, computationally designing effective bait sets is a challenging problem, which to the best of our knowledge, has not been formally defined or considered from an algorithmic perspective. Although there are a number of methods for designing baits—including MrBait [2], CATCH [13], and BaitFisher [12]—most methods are unable to scale or provide reasonable output to even moderate-sized sets of genomes.

We formalize this problem, which we call The `Minimum Bait Cover Problem`, and show it is NP-hard even for extremely restrictive versions of the problem. For example, the problem remains intractable even for an alphabet of size four (i.e., `A, T, C, G`), a single reference genome, a bait length that is logarithmic in the length of the reference genome, and Hamming distance of zero. In light of these hardness results, we provide an efficient heuristic based on the FM-index and demonstrate that it is capable of scaling to large sets of sequences. We refer to our method as syotti^*^ and compare it to all competing methods on three datasets: (1) MEGARes, which is a database of antimicrobial resistance (AMR) genes, (2) a set of genomes corresponding to substrains of Salmonella enterica subsp. enterica, Escherichia coli, Enterococcus spp. and Campylobacter jejuni, and (3) a set of viral genomes. We determined that syotti is the only method capable of scaling to the second and third datasets. MrBait was able to process all datasets, but was over 20 times slower and produced over 40 times more baits than syotti, making it unreasonable in practice. CATCH failed to process both the second and third dataset in 24 hours, and even failed to process prefixes of length 8% and 5% respectively in the same time limit. The curve of the running time of CATCH indicates that it would take at least 20 days to process the viral dataset, and probably much longer.

Lastly, we evaluated the coverage of the genomes with respect to the bait sets using our implementation of the evaluation method provided with CATCH [13]. On the smallest dataset (MEGARes), both syotti and CATCH covered 100.0% of the nucleotide positions at least once, but MrBait covered only 96.4%. On the largest subset of bacteria genomes that all tools were able to run, syotti produced 158 thousand baits, CATCH produced 241 thousand, and MrBait over 1 million; hence, the bait set of MrBait was too large to be deemed of any practical use. Lastly, we considered the full set of viral genomes and compared the baits of syotti against the published and publicly available bait sets of size 250k, 350k and 700k published with CATCH. The coverage of the CATCH bait sets were 84.1%, 90.6% and 97.5% respectively. The coverage of syotti was 99.5%, with the remaining 0.5% being due to unknown N-characters in the data.

## 2 The Minimum Bait Cover Problem

### 2.1 Preliminaries

We define a string *S* as a finite sequence of characters *S* = *S*[1 … *n*] over an alphabet Σ = {*c*_1_, …, *c_σ_*}. We denote the length of a string *S* by |*S*|, and the empty string by *ε*. We denote by *S*[1 … *i*] the *i*-th prefix of *S*, and by *S*[i … *n*] the *i*-th suffix of *S*. Given two strings *S* and *T*, we say that *S* is lexicographically smaller than *T* if either *S* is a prefix of *T* or there exists an index *i* ≥ 1 such that *S*[1 … *i*] = *T* [1 … *i*] and *S*[*i* + 1] occurs before *T* [*i* + 1] in Σ. We denote this as *S ≺ T*. Next, we define the *Hamming distance* between *S* and *T* (which have the same length) as the number of positions where *S* and *T* mismatch, namely *d*(*S, T*) ≔ |{*i* ≠ *S*[*i*] ≠ *T* [*i*]}|. Lastly, we denote *S* ○ *T* for the string concatenation of *S* and *T*. Given a set of strings, 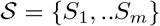, we denote the total length 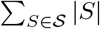 of all strings in 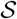 by 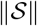.

For a string *S* and integer *ℓ* ≤ |*S*| we denote by pre(*S, ℓ*) the prefix *S*[1, …, *ℓ*] of *S* of length *ℓ*. Similarly we denote by suf(*S, ℓ*) the suffix *S*[*ℓ* − *ℓ* + 1, …, *ℓ*] of *S* of length *ℓ*.

### 2.2 Hardness Results

We fix the length of the bait sequences to a constant *L*. We say a string *X covers* a position *i* ≤ |*Y*| in a string *Y* if there exists a *j* such that *i* − |*X*| < *j* ≤ *i* and *X* = *Y* [*j, j* + 1, … *j* + |*X*| − 1]. More generally, *X distance θ-covers* (we will just say *θ*-*covers*) position *i* in *Y* if there exists a *j* such that *i* − |*X*| < *j* ≤ *i* and *d*(*X, Y* [*j, j* + 1, … *j* + |*X*| − 1]) ≤ *θ*. Observe that *X* covers *i* if and only if *X* 0-covers *i*. A set 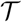 of strings *θ*-cover a string *S* if, for every position *i* in *S* there exists a string 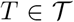 that *θ*-covers *i*. A set 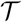 of strings *θ*-cover a set 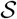 of strings if every string 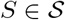 is *θ*-covered by 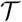. We are now ready to define the main problem considered in this paper, namely Minimum Bait Cover.

##### Minimum Bait Cover

Input: Here the input consists of two integer parameters *θ* ≥ 0 and *L* > 0 and a set 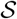 of *n* strings 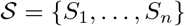 over a finite alphabet Σ.

Question: What is the smallest possible set 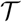 of strings, each of length exactly *L*, such that 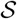 is *θ*-covered by 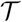.

In the closely related String Cover problem input is a single string *S* and a parameter *L* > 0. The task is to find a minimum size set 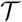 of strings, each of length exactly *L*, such that *S* is *θ*-covered by 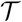. Comparing the two problem definitions it is easy to see that String Cover is precisely equal to the special case of Minimum Bait Cover when *n* = 1 and *θ* = 0. Cole et al. [3] proved that String Cover is `NP`-hard even for *L* = 2. This immediately leads for the following hardness result for Minimum Bait Cover.

#### Proposition 2.1.

*[3] (a)* String Cover *is `NP`-hard even for L* = 2*. (b)* Minimum Bait Cover *is `NP`-hard, even for n* = 1, *θ* = 0 *and L* = 2.

Proposition2.1 effectively rules out any hope of a polynomial time algorithm, or an efficient (Fixed Parameter Tractable or Slicewise Polynomial) parameterized algorithm with parameters *n*, *θ*, and *L*. At the same time the instances of String Cover constructed in the reduction of Cole et al. [3] are strings over a large alphabet |Σ|. Since we are primarily interested in instances of Minimum Bait Cover with |Σ| = 4, this might leave some hope that for small alphabets one can still obtain efficient parameterized algorithms. We now show that (almost) all of the hardness from Proposition 2.1 is retained even when |Σ| is constant.

#### Theorem 2.2.

(a) *For every k* ≥ 2, String Cover *is `NP`-hard with* |Σ| = *k and L* = *O*(log |*S*|). (b) *For every k* ≥ 2, Minimum Bait Cover *is `NP`-hard, even for* |Σ| = *k,* 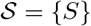, *θ* = 0 *and L* = *O*(log |*S*|).

Theorem 2.2 shows that unless `P` = `NP` there can’t exist an algorithm for Minimum Bait Cover with running time 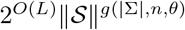 for any function *g* of constant parameters |Σ|, *n* and *θ*. This effectively rules out parameterized algorithms that exploit any of the most natural parameters one could expect to be small in relevant input instances. We remark that part (a) of Theorem 2.2 resolves in the affirmative a conjecture of Cole et al. [3] that String Cover remains `NP`-hard even with a constant size alphabet.

*Proof of Theorem 2.2*. We only prove (*a*), since (*b*) follows from (*a*) together with the fact that String Cover is the special case of Minimum Bait Cover with *θ* = 0 and *n* = 1. Fix an integer *k* ≥ 2. We give a reduction from String Cover with *L* = 2 and unbounded alphabet size (i.e |Σ| ≤ |*S*|) to String Cover with alphabet Σ_*k*_ = {0, …, *k* − 1} of size exactly *k*.

We now describe the construction. Given as input an instance (Σ*, S, L* = 2) of String Cover the reduction algorithm sets Σ_*k*_ = {0, …, *k* − 1}, *ℓ* to be the smallest integer above ⌊100 log_2_(|Σ|)⌋ that is divisible by 3, and *L*′ = 2*ℓ*. The algorithm then computes a set of strings {*S_a_* : *a* ∈ Σ} by using Claim 2.3.

#### Claim 2.3.

*There exists an algorithm that given as input a set* Σ, *runs in time polynomial in* |Σ| *, and outputs a set {S_a_* : *a* ∈ Σ} *of strings over the alphabet* Σ_*k*_, *all of length L*′ = 2*ℓ, that satisfy the following properties*:

1. *For distinct characters a, b* ∈ Σ *we have `pre`*(*S_a_, ℓ/*3) ≠ `pre`(*S_b_, ℓ/*3),
2. *for distinct characters a, b* ∈ Σ *we have `suf`*(*S_a_, ℓ/*3) = `suf`(*S_b_, ℓ/*3),
3. *for every pair of characters a, b* ∈ Σ *(including the case when a* = *b) and every r* ≥ *ℓ/*3 *we have that `pre`*(*S_a_, r*) ≠ `suf`(*S_b_, r*).

*Proof.* Let us first observe that the total number of strings in 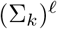 is 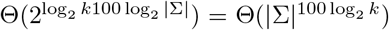 and therefore upper bounded by a polynomial in |Σ|. The algorithm initializes a list of strings 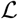 to contain all strings in (Σ_*k*_)^*ℓ*^. The algorithm then removes from 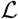 all strings *T* such that there exists some *r* ≥ *ℓ/*3 such that `pre`(*T, r*) ≠ `suf`(*T, r*). For every *r* ≥ *ℓ/*3, the number of strings *T* ∈ (Σ_*k*_)^*ℓ*^ such that the prefix of *T* of length *r* is equal to the suffix of *T* of length *r* is at most 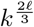 (because fixing the 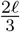 first characters uniquely determines the remaining 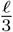 characters). Thus, the total number of strings removed from the list in this initial cleaning step is at most 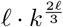.

The algorithm then iterates over the characters *a* ∈ Σ one by one. When considering the character *a* the algorithm picks an arbitrary string still on the list 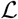, calls it *S_a_*, and removes it from the list 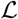. It then goes over all strings on the list 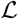 and removes all strings *T* such that `pre`(*T, ℓ/*3) = `pre`(*S_a_, ℓ/*3), or `suf`(*T, ℓ/*3) = `suf`(*S_a_, ℓ/*3), or there exists an *r* ≥ *ℓ/*3 such that `pre`(*T, r*) = `suf`(*S_a_, r*) or `suf`(*T, r*) = `pre`(*S_a_, r*).

There are at most 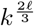 strings *T* such that `pre`(*T, ℓ/*3) = `pre`(*S_a_, ℓ/*3) (since 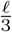 of the characters of *S* are uniquely determined by `pre`(*S_a_, ℓ/*3)). Similarly, there are at most 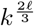 strings *T* such that `suf`(*T, ℓ/*3) = `suf`(*S_a_, ℓ/*3), and at most 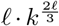 strings *T* such that there exists an *r ≥ ℓ/*3 such that `pre`(*T, r*) = `suf`(*S_a_, r*) or `suf`(*T, r*) = `pre`(*S_a_, r*). Therefore, in each iteration (i.e after selecting one string *S_a_*) the algorithm removes at most 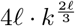 strings from 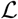

There are 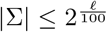 iterations of the algorithm in total. Thus, in each iteration of the algorithm there are at least 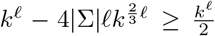 strings from 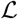 to choose the next string *S_a_* from. Thus, the algorithm successfully selects a string 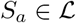 for every character *a* ∈ Σ.

Finally we argue that the set of strings {*S_a_* : *a* ∈ Σ} satisfy properties 1, 2 and 3. Suppose for distinct characters *a*, *b* we have that `pre`(*S_a_, ℓ/*3) = `pre`(*S_b_, ℓ/*3) or `suf`(*S_a_, ℓ/*3) = `suf`(*S_b_, ℓ/*3), or there exists an *r* ≥ *ℓ/*3 such that `pre`(*S_a_, r*) = `suf`(*S_b_, r*) or `suf`(*S_a_, r*) = `pre`(*S_b_, r*). Without loss of generality *S_a_* was selected before *S_b_*, and then *S_b_* would have been removed from 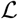 in the cleaning step immediately after *S_a_* was selected. This contradicts that *S_b_* was selected from the list 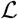. Further, suppose that there exists an *r* ≥ *ℓ/*3 such that `pre`(*S_a_, r*) = `suf`(*S_a_, r*) for some string *S_a_*. But then *S_a_* would have been removed in the initial cleaning step, again contradicting that *S_a_* was selected from 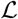. This completes the proof of the claim.

The reduction algorithm now constructs the string *S*′ from *S* by replacing every character *a* ∈ Σ with the corresponding string *S_a_*. The reduction algorithm outputs the instance (Σ′, *S*′, *L*′). This concludes the construction. We claim that for every integer *t*, there exists a set 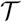 of *t* strings of length *L* = 2 that cover *S* if and only if there exists a set 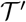 of *t* strings of length *L*′ that cover *S*′.

We begin by proving the forward direction of the above claim. Let 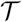 be a set of *t* strings of length *L* that cover *S*. We define 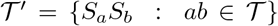. Clearly 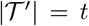. To see that 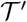 covers *S*′ consider an arbitrary position *p*′ in the string *S*′ and let 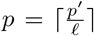. Since 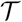 covers *p* there is a string 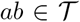 that covers *p* in *S*. But then 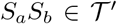 covers all positions {*ℓ · p* − *ℓ* + 1*, ℓ · p − *ℓ* + 2*, *ℓ* · *p*} in *S*′. However *p*′ ∈ {*ℓ · p* − + 1*, ℓ · p* − *ℓ* + 2*, ℓ · p* and therefore 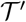 covers *p*′. Since *p*′ was an abitrarily chosen position we conclude that 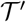 covers *S*′.

We now prove the reverse direction: if there exists a set 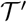 of *t* strings of length *L*′ that cover *S*′ then there exists a set 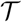 of *t* strings of length *L* = 2 that cover *S*. Without loss of generality, every string 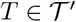 is a substring of *S*′ of length 2*ℓ* (otherwise 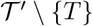 also covers *S*′). Thus *T* = `suf`(*S_a_, i*) ○ *S_b_* ○ `pre`(*S_c_*, *ℓ* − *i*) for some *i* ∈ {1, …, *ℓ*} and *a, b, c* ∈ Σ. Note that if *i* = *ℓ* then *T* = *S_a_S_b_* for *a, b* ∈ Σ. Next we prove a claim about strings on this form.

#### Claim 2.4.

*If `suf`*(*S_a_, i*) ○ *S_b_* ○ `pre`(*S_c_, ℓ* − *i*) = `suf`(*S_a′_, i*′)*S_b′_ `pre`*(*S_c′_, ℓ* − *i*′) *then i* = *i*′ *and b* = *b. Furthermore, if i* ≥ *ℓ/*2 *then a* = *a*′*, otherwise c* = *c*′.

*Proof.* Suppose that `suf`(*S_a_, i*) ○ *S_b_* ○ `pre`(*S_c_, ℓ* − *i*) = `suf`(*S_a′_, i*′)*S_b′_* `pre`(*S_c′_, ℓ* − *i*′), and assume for contradiction that *i* ≠ *i*′. Without loss of generality we have *i* < *i*′.

We have that: `suf`(*S_a′_, i*′ − *i*) = `pre`(*S_b_, i*′ − *i*), `suf`(*S_b_, ℓ* − *i*′ + *i*) = `pre`(*S_b′_, ℓ* − *i*′ + *i*), and `suf`(*S_b′_, i*′ − *i*) = `pre`(*S_c_, i* − *i*). Furthermore we have that |`suf`(*S_a′_, i*′ − *i*)| + |`suf`(*S_b_, ℓ* − *i*′ + *i*)| + |`suf`(*S_b′_, i*′ − *i*)| = *ℓ* + *i*′ − *i* ≥ *f*. Thus, at least one of `suf`(*S_a′_, i*′ − *i*), `suf`(*S_b_, ℓ* − *i*′ + *i*), or `suf`(*S_b′_, i*′ − *i*) has length at least *ℓ/*3. However, either one of `suf`(*S_a′_, i*′ − *i*), `suf`(*S_b_, ℓ* − *i*′ + *i*), or `suf`(*S_b′_, i*′ − *i*) having length at least *ℓ/*3 contradicts Property 3 of the set {*S_a_* : *a* ∈ Σ} of strings. We conclude that *i* = *i*′. But then `suf`(*S_a_, i*) = `suf`(*S_a′_, i*), *S_b_* = *S_b′_* and `pre`(*S_c_, ℓ* − *i*) = `pre`(*S_c′_, ℓ* − *i*). Property 1 implies that *b* = *b*′. If *i* ≥ *ℓ/*2 then Property 2 implies that *a* = *a*′. If *i* < *ℓ/*2 then *ℓ* − *i* ≥ *ℓ/*3 and therefore Property 1 implies that *c* = *c*′.

For every string 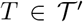 we have that *T* = `suf`(*S_a_, i*) ○ *S_b_* ○ `pre`(*S_c_, ℓ* − *i*) for some *i* ∈ {1, …, *ℓ*} and *a, b, c* ∈ Σ. By Claim 2.4 *i* and *b* are uniquely determined by *T*. If *i* ≥ *n/*2 then by Claim 2.4 *a* is also uniquely determined by *T*. In this case we add *ab* to the set 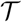. If *i* < *ℓ/*2 then by Claim 2.4 *c* is also uniquely determined by *T*. In that case we add *bc* to the set 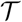. In either case we add precisely one string of length 2 to 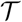 for each 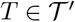. Thus 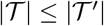 and it remains to prove that 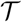 covers *S*.

Consider an arbitrary position *p* of *S* and let *abcde* = *S*[*p* − 2, *p* − 1*, p, p* + 1*, p* + 2]. Let 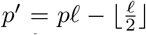, and let 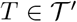 be such that *T* = *S*[*x, x* + 1*, …, x* + *L*′ − 1] where *p*′ ∈ {*x, x* + 1*, …, x* + *L*′ − 1}. We have the following cases:

1. *T* = `suf`(*S_a_, i*) ○ *S_b_* ○ `pre`(*S_c_, ℓ* − *i*), where *ℓ* − *i* ≥ *ℓ/*2, or
2. *T* = `suf`(*S_b_, i*) ○ *S_c_* ○ `pre`(*S_d_, ℓ* − *i*), where *i* ∈ 1, …, *ℓ*, or
3. *T* = `suf`(*S_c_, i*) ○ *S_d_* ○ `pre`(*S_e_, ℓ* − *i*), where *i* ≥ *ℓ/*2.

In the first case 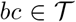 and *S*[*p* − 1*, p*] = *bc*, in the second case *bc* ∈ *T* and *S*[*p* − 1*, p*] = *bc* or *cd* ∈ *T* and *S*[*p, p* + 1] = *cd*, while in the third case 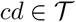 and *S*[*p, p* + 1] = *cd*. In either case the position *p* is covered by 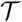, and so *S* is covered by 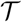.

We remark that in the argument above, if *p* ∈ {1, 2, |*S*| − 1, |*S*|} then some of *a*,*b*,*d*,*e* are not properly defined. However this only restricts which of the cases 1, 2 and 3 may apply, one of them must still apply (for characters from *a*,*b*,*c*,*d*,*e* that are well defined). This concludes the proof.

**Algorithm 1.**
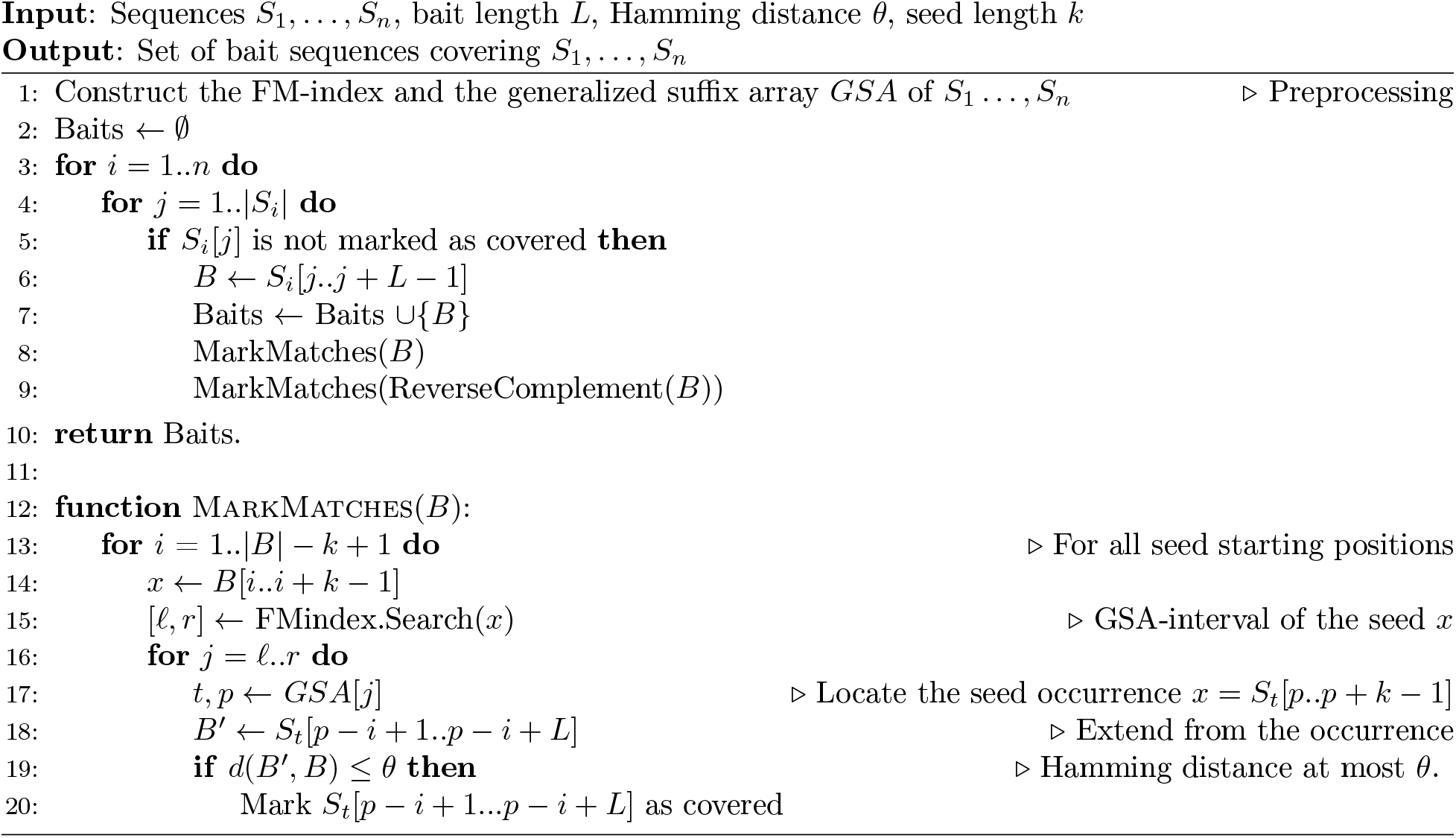
The syotti algorithm.

### 2.3 The Syotti Algorithm

Our algorithm requires an index that can search for substrings in the input sequences *S*_1_, …, *S_n_* and report the locations of the exact matches. For this purpose, we use the combination of an FM-index [6] and a generalized suffix array *GSA* (the suffix array of a set of strings [18]). Given a substring *x*, the FM-index is able to compute an interval [*ℓ, r*] such that the entries *GSA*[*ℓ*], …, *GSA*[*r*] give all starting positions of *x* in *S*_1_, …, *S_n_*. That is, the entries of *GSA* in the interval are pairs (*t, p*) such that *S_t_*[*p..p* + |*x*| − 1] = *x*. We refer the interested reader to Mäkinen et al. [11] for a detailed exposition of the techniques involved.

First, we construct the aforementioned data structures from the input sequences. Then, we initialize the set of bait sequences to an empty set, and initialize a bit vector for each sequence, of the same length as that sequence. The bit vector signifies which positions are covered by the bait sequences. Next, we perform a linear scan of each sequence, checking the bit vector at each position. When we come across a position that is not yet marked as covered, we add the substring of length *L* starting from that position into the set of bait sequences and update the bit vectors for all positions covered by the new bait. More specifically, we search for positions in all the sequences to which the new bait has Hamming distance less than or equal to *θ*. This search is done with a seed-and-extend heuristic based on the exact matching index as follows. For each seed *k*-mer *x* in the new bait sequence *B*, we find all occurrences of *x* in our input sequences. For each occurrence of *x* we consider the *L*-length sequence *B*′ that contains *x* in the same position as in *B*. If the Hamming distance of *B* and *B*′ is at most *θ*, we update the bit vector; otherwise, we move onto the next occurrence of *x*. After processing all seeds of *B*, we repeat the process for the reverse complement of *B*.

We give the pseudocode in Algorithm 1. There are some implementation details omitted from the pseudocode to avoid obfuscating the main idea of the algorithm. The string indices can go past the ends of the strings, but these corner cases are handled easily. That is, if a bait would run past the end of the string, we shift it back inside the string, and we ignore candidate matches from seeds that run past either end of the string. Also, if implemented directly as in the pseudocode, the seed-and-extend algorithm often ends up comparing the bait against the same position from multiple seeds in the same bait. To avoid this, we store the starting positions of the candidate matches in a set data structure that keeps only the distinct starting positions. We then run the Hamming distance computation on each distinct candidate match.

## 3 Experiments

We implemented syotti in the C++ programming language. We use the Divsufsort and SDSL libraries [8] for the construction of the FM-index and the generalized suffix array required by the syotti algorithm. The implementation of syotti is publicly available at https://github.com/jnalanko/syotti.

The main competitor for syotti is the method CATCH by Metsky et al. [13]. CATCH reduces the bait design problem to an instance of the classic NP-hard set cover problem, and finds an approximate solution by greedily picking baits in decreasing order of profitability. Two other potential competitors are MrBait [2] and BaitFisher [12]. MrBait employs a bait tiling strategy followed by optional post-processing filtering, and BaitFisher works by clustering sequences using a multiple alignment and tiling the consensus sequences of the clusters with baits.

We evaluate our method and the competing methods on three datasets: (1) `MEGARES`, which is a database containing 7,868 AMR genes of total length 8,106,325 bp, publicly available at the MEGARes website: https://megares.meglab.org/download. (2) `VIRAL`, which is a set of 422,568 viral genomes from CATCH [13], of total length 1,257,789,768 bp, publicly available at: https://github.com/broadinstitute/catch. (3) `BACTERIA`, which is a custom (see Appendix A) set of 1,000 bacterial genomes representing four foodborne pathogens containing 98,984 sequences of total length 3,040,260,476 bp.

As advised by the bait manufacturer Agilent, we set the bait length to 120 and tolerated 40 mismatches. The seed length in syotti was set to 20. We ran CATCH, MrBait, and syotti on these datasets on a server with two Intel Xeon Gold 5122 CPUs at 3.60GHz for total of 8 physical cores and up to 16 threads with hyperthreading, 503GB of available memory and a 3TB NVMe SSD for storage. The time is measured in wall-clock seconds. The experiments do not include BaitFisher because it requires a multiple sequence alignment of all the input sequences as input, but our sequences do not all align well to each other. Feeding multiple alignments computed with Clustal Omega [19] to BaitFisher produced extremely poor baits sets that had more nucleotides than the original sequences.

CATCH was run with the command line options `-pl 120 -m 40 -l 120 -ps 60`. We also experimented with the options `--cluster-and-design-separately 0.15` and `--cluster-from-fragments 10000`, but did not see significant improvement on the results or the run time. MrBait was run with the options `-A 120 -s tile=0`. MrBait also offers post-processing filtering using all-pairs global alignment from the external tool VSEARCH [16]. This was feasible only for `MEGARES` because the time complexity of all-pairs alignment is quadratic. syotti was run with options `-L 120 -d 40 -r -t 16 -g 20`.

We restricted all methods to using 24 hours of time, and the available 503 GB of memory, and 3 TB of disk space. None of the methods exceeded the memory and disk limits before being terminated by the 24 hour time limit. We analyzed the coverage of the produced bait sets by mapping the baits to the reference sequences using the seed-and-extend method with seed length 20, tolerating up to 40 mismatches. This is the same method of analysis as the one used in the coverage analyser of CATCH. We did not use the CATCH analyser because it does not scale to our larger datasets. To verify the comparability of the two approaches, we ran them both on small inputs and found negligible difference.

### 3.1 Results on Antimicrobial Resistance Genes

We ran all methods on exponentially increasing subsets of the MEGARes database that consists of 7,868 AMR genes. All methods were capable of being run on the complete dataset. The memory, running time, and number of baits is given in Table 1. The scaling of the time usage is plotted in Figure 1.

**Table 1:**
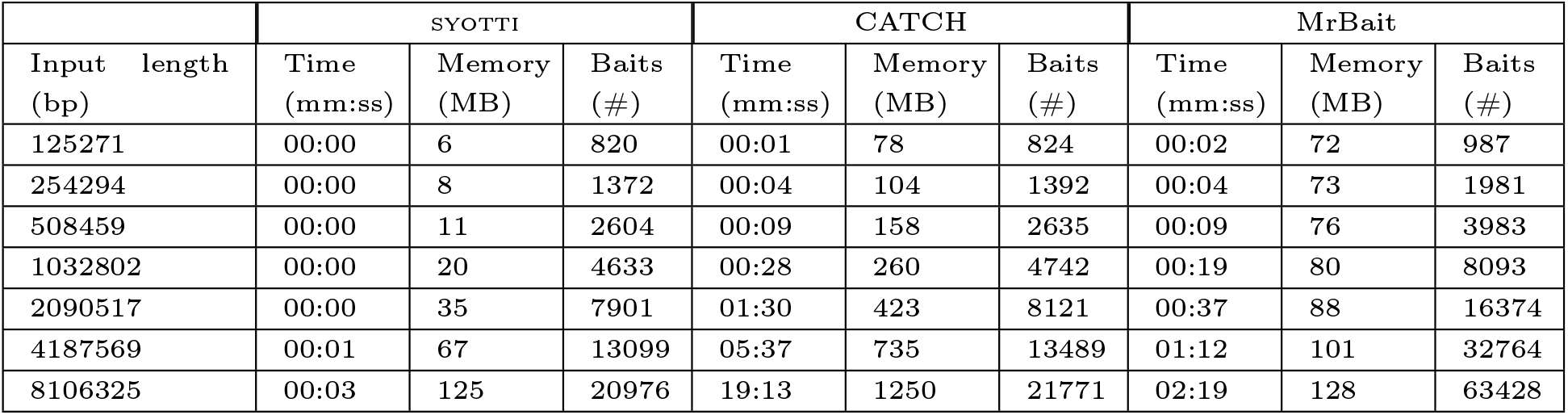
Results on the dataset `MEGARES` (without VSEARCH filtering). The seconds are rounded down. Rows where all tools ran in less than one second have been removed. The full table is in Appendix B.

|  | SYOTTI |  |  | CATCH |  |  | MrBait |  |  |
| --- | --- | --- | --- | --- | --- | --- | --- | --- | --- |
| Input length (bp) | Time (mm:ss) | Memory (MB) | Baits (#) | Time (mm:ss) | Memory (MB) | Baits (#) | Time (mm:ss) | Memory (MB) | Baits (#) |
| 125271 | 00:00 | 6 | 820 | 00:01 | 78 | 824 | 00:02 | 72 | 987 |
| 254294 | 00:00 | 8 | 1372 | 00:04 | 104 | 1392 | 00:04 | 73 | 1981 |
| 508459 | 00:00 | 11 | 2604 | 00:09 | 158 | 2635 | 00:09 | 76 | 3983 |
| 1032802 | 00:00 | 20 | 4633 | 00:28 | 260 | 4742 | 00:19 | 80 | 8093 |
| 2090517 | 00:00 | 35 | 7901 | 01:30 | 423 | 8121 | 00:37 | 88 | 16374 |
| 4187569 | 00:01 | 67 | 13099 | 05:37 | 735 | 13489 | 01:12 | 101 | 32764 |
| 8106325 | 00:03 | 125 | 20976 | 19:13 | 1250 | 21771 | 02:19 | 128 | 63428 |

**Figure 1:**
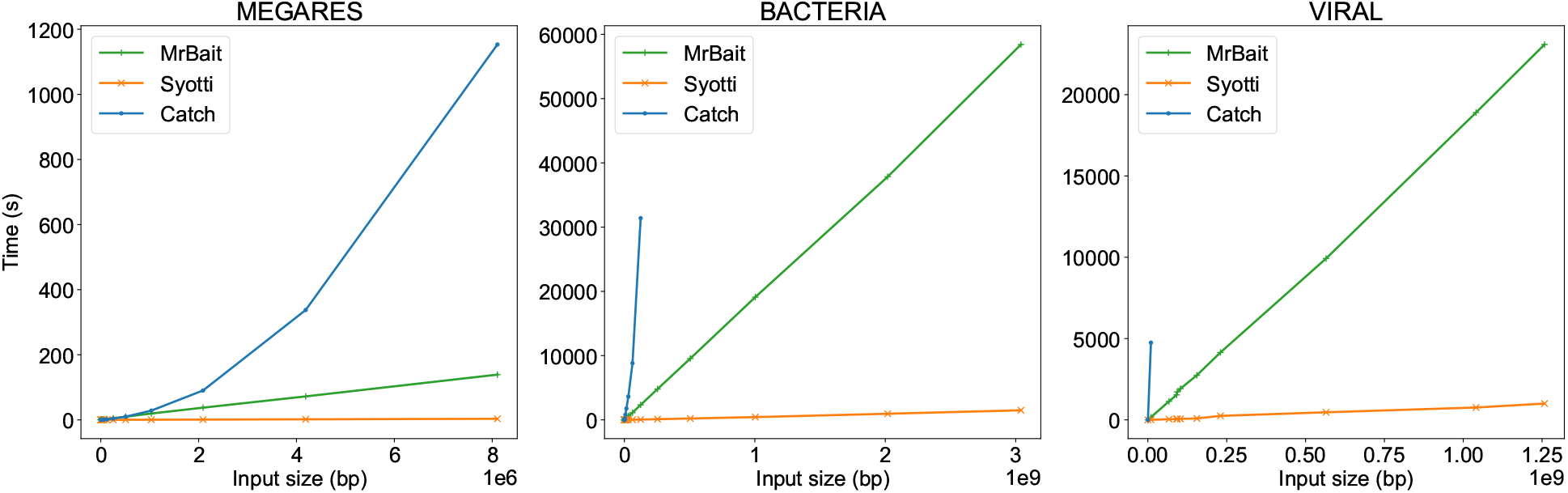
Time scaling on all three datasets.

Even though the worst case running time of syotti is at least quadratic in the length of the input, the running time behaved approximately linearly. MrBait also behaved approximately linearly, but up to 40 times slower than syotti. On the other hand, CATCH showed an upward-bending growth curve, which we believe is due to having to update the profitability scores of candidate baits after every iteration, resulting in a quadratic time complexity in practice as well as in the worst case.

The size of bait sets of CATCH and syotti had negligible difference, with the bait set of syotti being consistently slightly smaller. MrBait produced 2-3 times larger bait sets than the other two. The memory usage of syotti was the smallest of the three, however, the memory usage for all methods was less than 2 GB. The slope of the memory of MrBait was smaller than syotti, which indicates that the memory of syotti would grow past MrBait if the dataset size was still increased. We believe that MrBait’s good memory scaling is due to its use of a disk-based SQLite database.

Lastly, we evaluated the coverage of the baits. We note that the most optimal solution has a coverage value of 1, indicating sufficient likelihood of bait binding, but without redundancy (i.e., inefficiency) in the bait set. This is important for two reasons: first, multiple baits for the same position could create bait-bait interference during the binding process; and second, each bait is relatively costly, with a fixed upper limit on the total number of baits that can be manufactured. Coverage was analyzed by mapping the baits to the reference sequences with a seed-and-extend approach with a seed length of 20. The coverage of a position is defined as the number of mapped baits spanning that position with at most 40 mismatches. Figure 2 shows the mean coverage in each sequence of the database. Both syotti and CATCH covered 100.0% of the nucleotide positions at least once, but MrBait covered only 96.4%. The bait set designed by MrBait was extremely inefficient, with many positions being covered by hundreds of baits (Figure 2). The coverage efficiencies for CATCH and syotti were comparable, with a slight advantage for syotti (Figure 2). Filtering the MrBait baits with VSEARCH and adding the command line parameters `-f pw=0.9,0.9` to MrBait, resulted in 22,230 baits with coverage 95.9%, but the run time increased drastically from 2min 19s to 4h 22min 36s.

**Figure 2:**
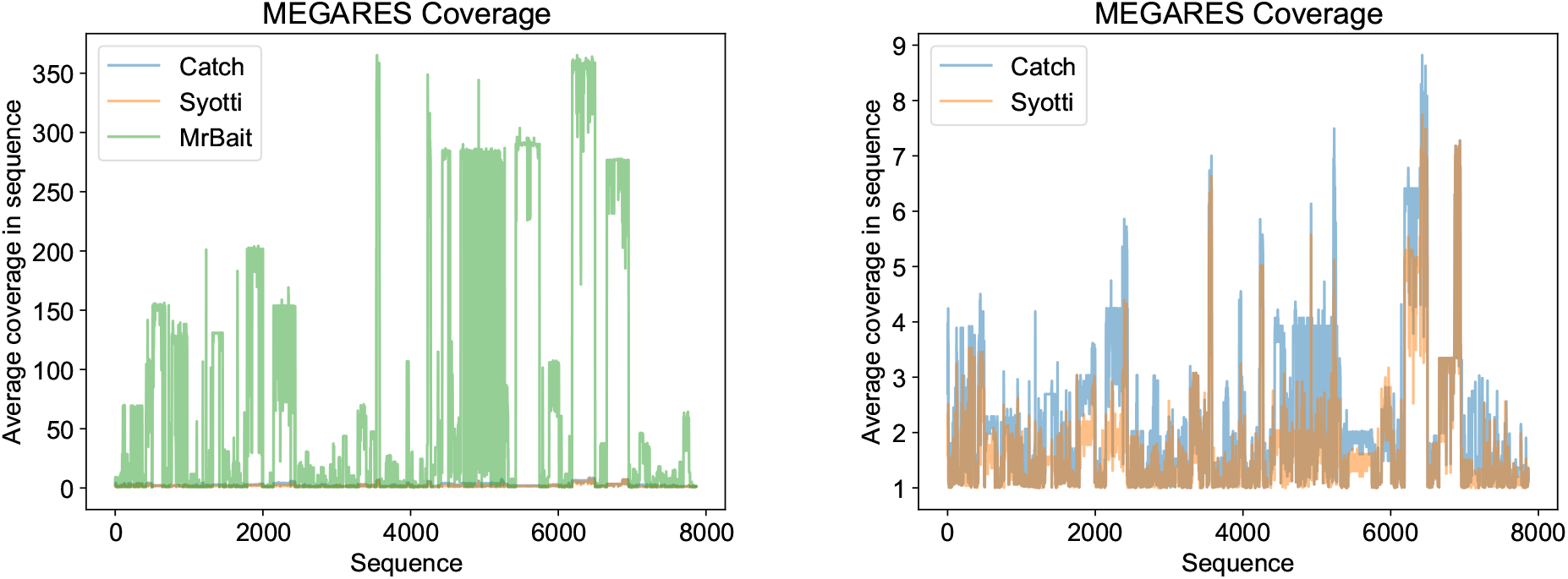
Average sequence coverage on the `MEGARES` dataset with all three methods (left) and with just CATCH and syotti (right). The sequences are in the order of the MEGARes database. The coverage plot with all tools after filtering MrBait baits with VSEARCH is in Appendix B.

### 3.2 Results on Bacterial Strains and Substrains

The aim of our second experiment was to study the scalability of the methods on pangenomes of clinically relevant bacterial species. Lacking a standard reference database of this type, we built our own, incorporating available sequences from foodborne pathogens *Salmonella enterica*, *Campylobacter jejuni*, *Escherichia coli*, *Enterococcus faecalis*. The purpose of the dataset is to cover as much of the known sequence diversity of these species as possible. The data was carefully selected and filtered to be suitable for the downstream application of foodborne pathogen detection. The full details of the process of compiling the dataset are available in Appendix A. We note that this database and bait set from syotti will be used in follow-up studies that aim to amplify these bacteria genomes in samples taken from food production facilities, in order to test for foodborne pathogens.

To measure the scalability of the methods, we ran them on subsets of the data containing 1,2,4,8,16 … 65536 sequences, as well as the full dataset consisting of all 98,984 sequences. The results are shown in Table 2. syotti and MrBait were able to process all inputs, but CATCH hit the 24 hour time limit on inputs larger than 250 million base pairs.

**Table 2:** Running time (Time), peak memory usage (Memory), and number of baits (baits) for increasingly larger subsets of sequences from the `BACTERIA` dataset. The times are in the format hh:mm:ss. “NA” signifies that the dataset surpassede 24 hours of running time. Rows where all tools took less than 30 minutes have been removed to save space. The full data table is in Appendix B

|  | SYOTTI |  |  | CATCH |  |  | MrBait |  |  |
| --- | --- | --- | --- | --- | --- | --- | --- | --- | --- |
| Input length | Time | Memory (MB) | Baits | Time | Memory (MB) | Baits | Time | Memory (MB) | Baits |
| 15920441 | 00:00:08 | 237 | 72035 | 00:29:32 | 4096 | 90182 | 00:04:48 | 195 | 132408 |
| 30786116 | 00:00:14 | 454 | 96595 | 01:00:27 | 7575 | 134112 | 00:10:00 | 316 | 256041 |
| 62502135 | 00:00:27 | 917 | 123541 | 02:26:56 | 14234 | 181825 | 00:18:44 | 572 | 519838 |
| 125063199 | 00:00:52 | 1831 | 157818 | 08:43:14 | 24269 | 240747 | 00:38:48 | 1076 | 1040174 |
| 254576853 | 00:01:45 | 3723 | 188813 | >24h | NA | NA | 01:20:03 | 2071 | 2117436 |
| 505422833 | 00:03:31 | 7387 | 223931 | NA | NA | NA | 02:39:03 | 4056 | 4203752 |
| 1003934029 | 00:07:11 | 14673 | 267890 | NA | NA | NA | 05:18:28 | 7980 | 8349888 |
| 2018459352 | 00:15:50 | 29496 | 324797 | NA | NA | NA | 10:30:37 | 15697 | 16788084 |
| 3040260476 | 00:24:52 | 44425 | 366761 | NA | NA | NA | 16:13:43 | 23615 | 25286576 |

**Table 3:**
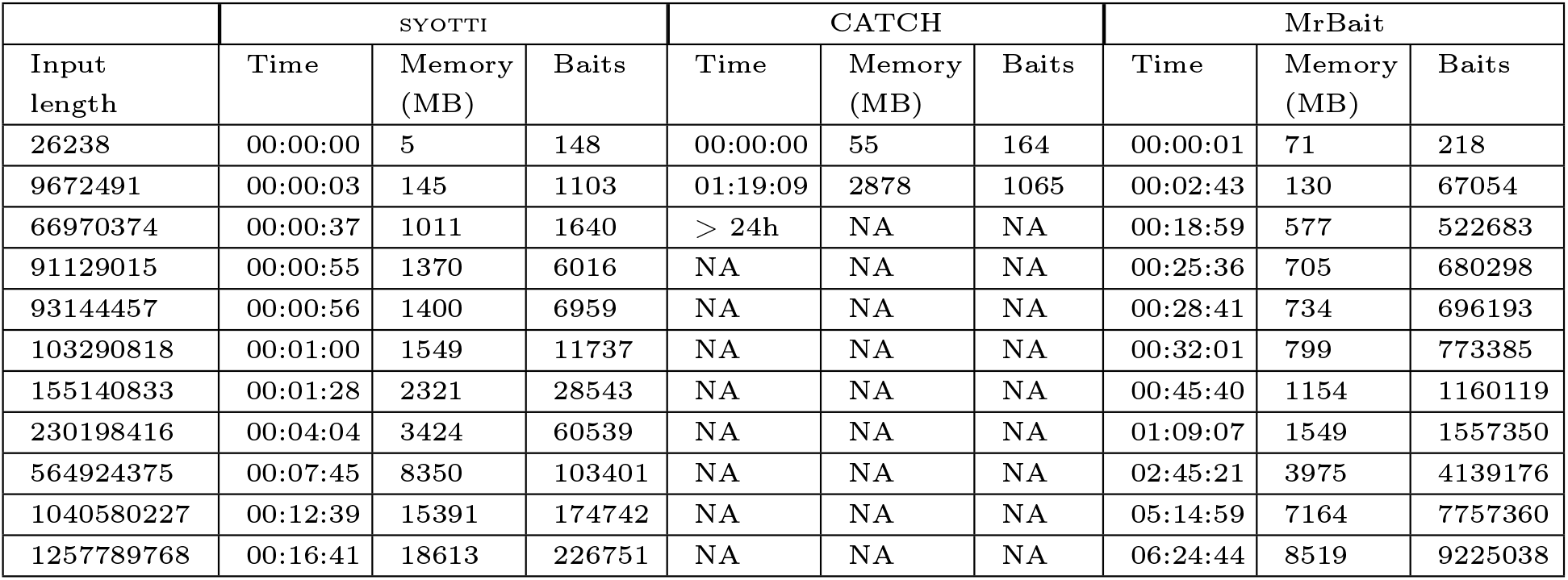
Running time (Time), peak memory usage (Memory), and number of baits (baits) for increasingly larger subsets of sequences from the `VIRAL` dataset. The times are in the format hh:mm:ss. “NA” signifies that the dataset surpassed 24 hours of running time.

All three tools produced bait sets of similar size for the smaller prefixes. But as the length of the prefix increased, the bait sets produced by syotti became significantly smaller than those produced by the other two tools. On the largest prefix that all tools were able to run, syotti produced 158 thousand baits, CATCH 241 thousand, and MrBait 1040 thousand. This showcases the ability of syotti to deal with multiple bacterial pangenomes at once. On the full dataset, syotti produced 367 thousand baits, whereas MrBait produced 25 million. Given the 220,000 limit provided by Agilent, two bait kits could be manufactured to include all our bait sequences, but over 113 bait kits would have to be manufactured for MrBait (which would cost more than 1 million USD for a single sample), making this bait set highly impractical.

The running times and memory usage of the tools showed similar trends to the first experiment on AMR sequences. syotti is consistently the fastest by an order of magnitude. MrBait was eventually the most efficient in terms of peak memory, thanks to its SQLite database on disk, but it pays a large price for the disk-based approach in the form of a significantly slower running time.

We observe that very good trade-offs between coverage and number of baits are available by halting the greedy algorithm of syotti before it reaches 100% coverage. Figure 4 shows the coverage as a function of the number of baits selected. For example, taking just the first 200k baits out of 367k already results in 98.8% coverage, whereas taking a *random* subset of 200k baits of the full set resulted in only 62.7% coverage. If long gaps in coverage are undesirable, we can add baits manually to patch the long gaps. For example, if we halt the greedy algorithm at 200k baits and then switch to filling the gaps such that the maximum gap length is 50 bp, we obtain a set of 275k baits with 99.6% coverage and with no gap longer than 50. Figure 4 shows the coverage plot for the 200k bait set after gap filling.

### 3.3 Results on Viral Strains and Substrains

For our final experiment, we downloaded the set of human viral pangenomes published in the Github repository of CATCH in 2018. We studied scaling of the tools by shuffling the list of files and ran the tools on prefixes containing 1, 2, 4, 8, 16*, …,* 512 files, as well as the whole dataset of 608 files.

syotti was again the fastest tool by an order of magnitude, producing 227k baits on the full dataset. MrBait again showed slow but approximately linear scaling, but the bait set was 40 times larger than the one produced by syotti. Surprisingly, CATCH itself was unable to process prefixes longer than 1% of the full dataset within the time limit of 24 hours. However, the Github page of CATCH contains pre-designed bait sets with 250k, 350k and 700k baits of length 75 for the dataset. The sets were designed by the authors of CATCH by running the tool separately for each species, and pooling the baits together with different parameters, optimizing the combined bait sets in the process.

To show CATCH in the best possible light, we compared the baits of syotti against the published and optimized bait sets of CATCH. We used syotti to generate baits of length 75, matching the bait length of the published data sets. The pooling method used with CATCH uses different values for the mismatch tolerance for different species to optimize the coverage. As syotti does not support a varying mismatch tolerance, the tolerance of syotti was set to 5, which is the maximum value in the parameters selected in Supplementary Table 1 of the manuscript of CATCH [13]. With these settings, syotti generated 684k baits.

We analyzed the coverage of the bait sets allowing up to 5 mismatches. Coverage of the CATCH bait sets of size 250k, 350k and 700k were 84.1%, 90.6% and 97.5% respectively. The coverage of syotti was 99.5%, with the remaining 0.5% being due to unknown N-characters in the data. The coverage plots of the bait set with 684k baits versus the CATCH bait sets are in Figure 3. Compared to the 700k baits of CATCH, syotti had mostly lower coverage than CATCH (i.e., higher efficiency), except for one notable stretch of the input which finished with 60-fold coverage, where CATCH was able to keep the coverage to only 10-fold. This stretch represents the 59,686 rotavirus A strains in the database. Taking the first 250k baits generated by syotti results in a coverage of 96.5%.

**Figure 3:**
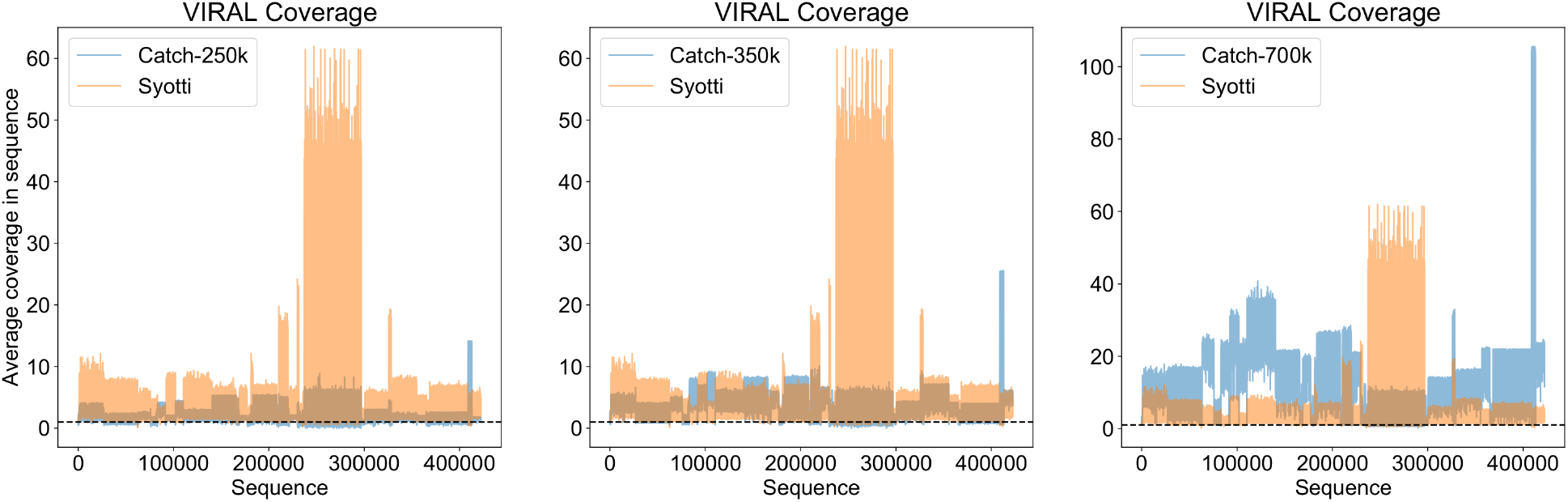
Coverage of the 684k baits generated by syotti for the `VIRAL` dataset, versus and the published bait sets of sizes 250k, 350k and 700k from CATCH. The dashed line shows coverage 1.0.

**Figure 4:**
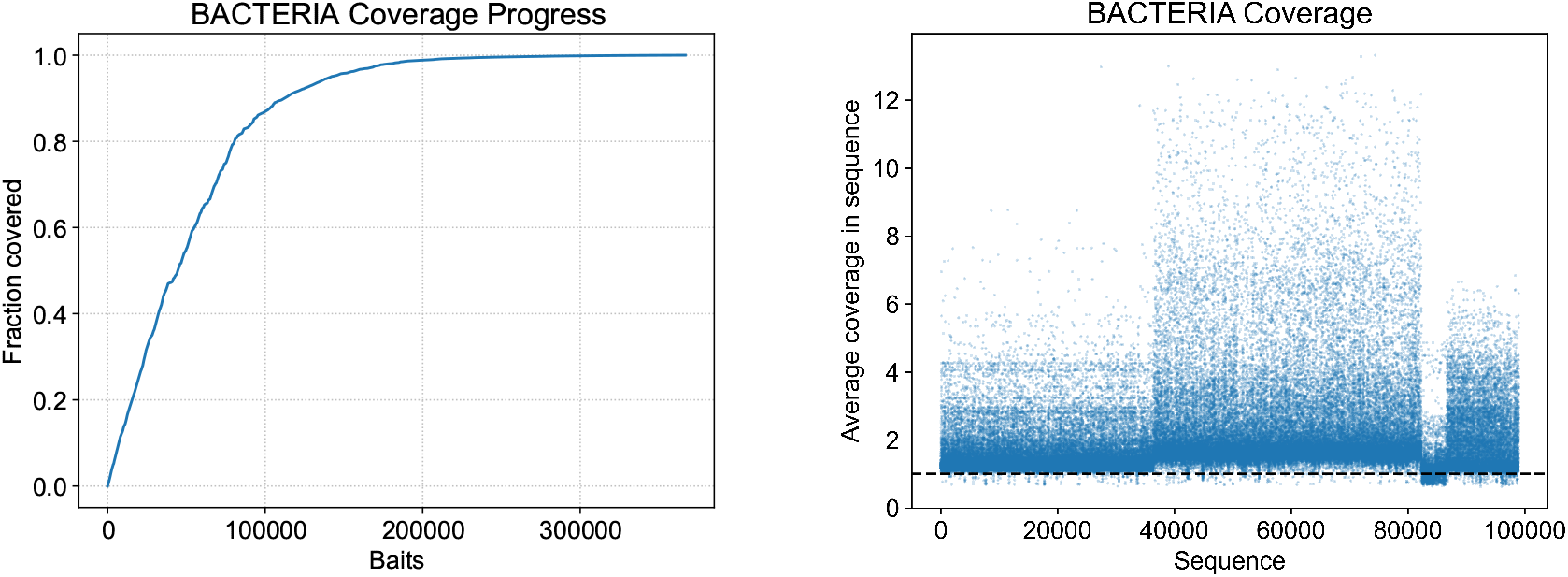
Left: Fraction of nucleotides covered by syotti as the algorithm progresses on the full `BACTERIA` dataset. Right: average coverage of the reference sequences in `BACTERIA` after taking the first 200k baits produced by syotti and filling gaps to maximum length 50. The dashed line shows coverage 1.0.

## 4 Conclusion

In this paper, we provide a formulation of designing baits and demonstrate that the problem is NP-hard even for deceptively simple instances, and provide a heuristic that works efficiently in practice. While both syotti and CATCH use a kind of greedy heuristic, the heuristic of syotti is designed to be much more efficiently implementable. We conclude by suggesting some areas that warrant future work. First, a more sophisticated binding model than the Hamming distance could be plugged into the heuristic at Line 19 of Algorithm 1. Second, while our results rule out any reasonable fixed parameter tractable algorithms, there is potential that the problem admits an approximation algorithm. However, we conjecture that the problem remains hard even for a constant approximation. Third, other artifacts of the laboratory process are left for consideration, including considering the GC-content of the baits; designing baits in a manner that avoids contaminants; and designing baits to avoid bait-to-bait binding. Lastly, computationally designing baits in a manner that models and controls off-target binding is worth consideration.

## A Details on how the Bacterial Dataset was Constructed

The pathogen set used was selected to represent whole genomes of public-health relevant foodborne isolates identified in food, animals, and human patients (clinical) using the NCBI Pathogen Detection tool [7], a repository developed for the curation of pathogenic bacterial genomic sequences. Salmonellae genomes were chosen to represent the serovars of greatest public health concern [1], including Enteriditis, Typhimurium, Infantis, Kentucky, Montevideo, and Dublin, representing an initial pool of 76,638 genomes and 3,486 SNP clusters. Shigatoxigenic Escherichia coli (STEC) isolates were chosen to include the O157:H7 serovars as well as the non-O157 STEC group (O26, O45, O103, O111, O121, and O145) including H-untypable, H-pending, H-undetermined, and non-motile types [5]. The resulting initial pool of STEC isolates represented 4,964 genomes and 1,164 SNP clusters. No specific subtype or serovar was chosen for Campylobacter jejuni (initial pool representing 60,227 genomes and 4,591 SNP clusters) or the Enterococci [20] (initial pool of 24,221 genomes and 1,774 SNP clusters of faecium and faecalis).

All isolates were then further filtered according to location (known U.S. and U.S. territories only), source (either from a clinical sample or from a domesticated animal or fowl collected pre- or post-slaughter, pre- or post-processing, and / or pre- or immediately post-fabrication stage in the food production system), collection date (2018-2020), sequence identity (i.e., availability of isolate-level WGS accession), and availability of SNP cluster assignment. Isolates were excluded if they originated from environmental samples (e.g., soil, water, sewage), ready-to-eat (RTE) products, or if any metadata variables were designated as ‘unknown’, ‘pending’, or left blank.

Stratified random sampling was conducted to select 1,000 total isolate genomes representative of the relative proportions of genera and serotypes in the final filtered pool of isolates (Salmonella enterica ssp. enterica: 322; STEC: 60; Campylobacter jejuni: 580; and Enterococci: 42). Corresponding WGS assemblies for all 1,000 isolates containing either a GCA or GCF prefix were retrieved from NCBI using the EDirect tool, and integrity of transferred data was ensured by evaluating file md5 checksum values.

## B Full Experimental Data Tables and Figures

All coverge plots, all time/mem plots. Full tables in case we delete some rows corresponding to the very smallest data prefixes.

**Table 4:** Running time, memory usage and number of baits for increasing larger subsets of the `MEGARES` dataset. Times are in the format mm:ss.

|  | SYOTTI |  |  | CATCH |  |  | MrBait |  |  |
| --- | --- | --- | --- | --- | --- | --- | --- | --- | --- |
| Input length | Time | Memory (MB) | Baits | Time | Memory (MB) | Baits | Time | Memory (MB) | Baits |
| 498 | 00:00 | 5 | 5 | 00:00 | 49 | 5 | 00:01 | 71 | 4 |
| 1320 | 00:00 | 5 | 12 | 00:00 | 49 | 12 | 00:00 | 71 | 10 |
| 2937 | 00:00 | 5 | 26 | 00:00 | 50 | 26 | 00:00 | 71 | 22 |
| 6672 | 00:00 | 5 | 59 | 00:00 | 50 | 59 | 00:00 | 71 | 52 |
| 14244 | 00:00 | 5 | 115 | 00:00 | 52 | 118 | 00:00 | 71 | 111 |
| 31758 | 00:00 | 5 | 249 | 00:00 | 56 | 253 | 00:01 | 71 | 249 |
| 60670 | 00:00 | 5 | 434 | 00:00 | 63 | 437 | 00:01 | 71 | 477 |
| 125271 | 00:00 | 6 | 820 | 00:01 | 78 | 824 | 00:02 | 72 | 987 |
| 254294 | 00:00 | 8 | 1372 | 00:04 | 104 | 1392 | 00:04 | 73 | 1981 |
| 508459 | 00:00 | 11 | 2604 | 00:09 | 158 | 2635 | 00:09 | 76 | 3983 |
| 1032802 | 00:00 | 20 | 4633 | 00:28 | 260 | 4742 | 00:19 | 80 | 8093 |
| 2090517 | 00:00 | 35 | 7901 | 01:30 | 423 | 8121 | 00:37 | 88 | 16374 |
| 4187569 | 00:01 | 67 | 13099 | 05:37 | 735 | 13489 | 01:12 | 101 | 32764 |
| 8106325 | 00:03 | 125 | 20976 | 19:13 | 1250 | 21771 | 02:19 | 128 | 63428 |

**Figure 5:**
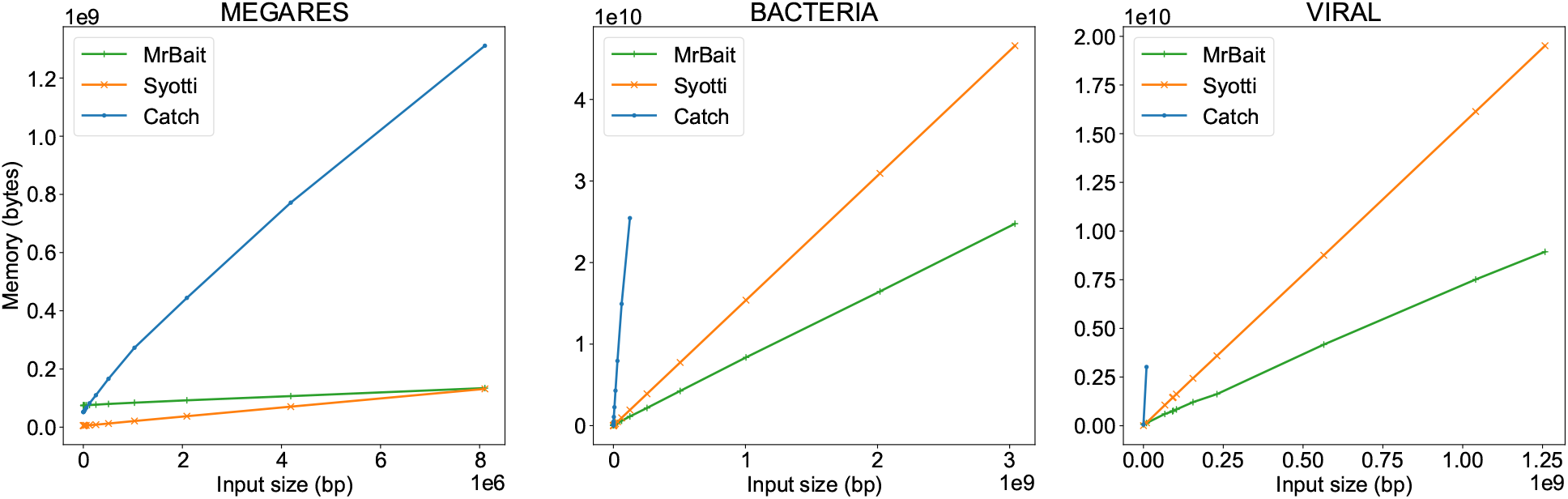
Memory scaling on all three datasets.

**Figure 6:**
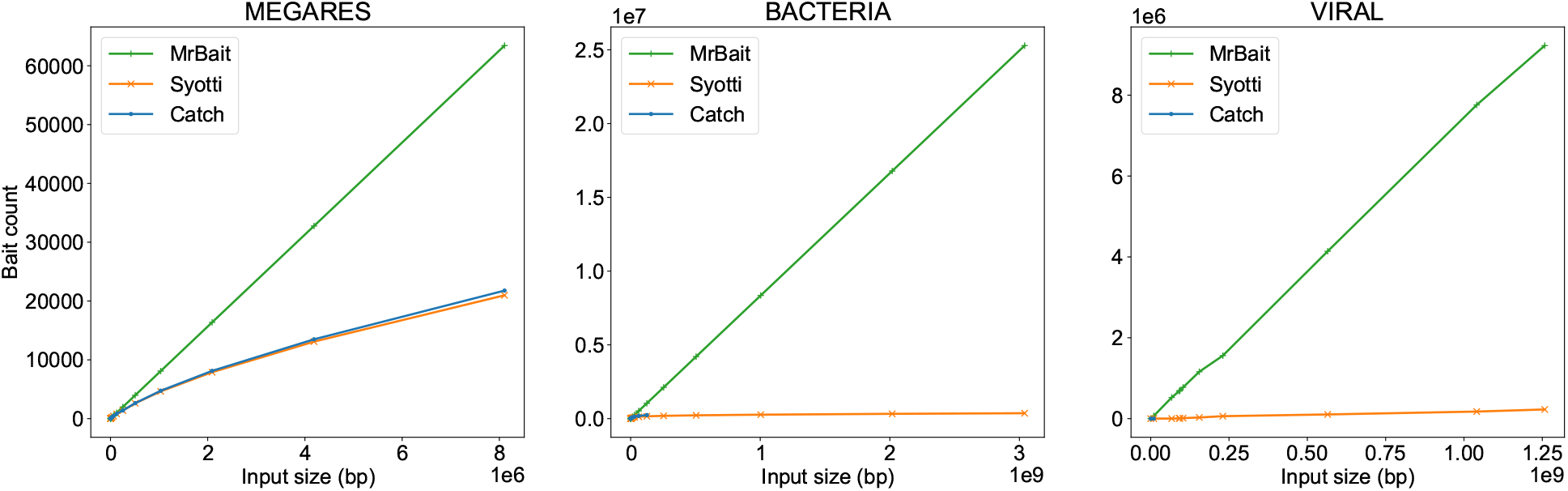
Bait set size scaling on all three datasets.

**Table 5:** Running time, memory usage and number of baits for increasing larger subsets of the `BACTERIA` dataset. The times are in the format hh:mm:ss.

|  | SYOTTI |  |  | CATCH |  |  | MrBait |  |  |
| --- | --- | --- | --- | --- | --- | --- | --- | --- | --- |
| Input length | Time | Memory (MB) | Baits | Time | Memory (MB) | Baits | Time | Memory (MB) | Baits |
| 16794 | 00:00:00 | 5 | 140 | 00:00:00 | 52 | 140 | 00:00:01 | 71 | 139 |
| 27058 | 00:00:00 | 5 | 226 | 00:00:00 | 55 | 226 | 00:00:00 | 71 | 224 |
| 164612 | 00:00:00 | 7 | 1373 | 00:00:02 | 95 | 1373 | 00:00:03 | 72 | 1369 |
| 242069 | 00:00:00 | 7 | 2017 | 00:00:03 | 112 | 2017 | 00:00:05 | 73 | 2012 |
| 344651 | 00:00:00 | 9 | 2875 | 00:00:05 | 146 | 2875 | 00:00:06 | 74 | 2862 |
| 991610 | 00:00:00 | 19 | 8189 | 00:00:29 | 312 | 8264 | 00:00:19 | 80 | 8244 |
| 1572067 | 00:00:00 | 27 | 12821 | 00:01:01 | 485 | 12975 | 00:00:30 | 84 | 13067 |
| 3470929 | 00:00:02 | 55 | 25351 | 00:03:47 | 1022 | 27378 | 00:01:07 | 98 | 28857 |
| 8038466 | 00:00:04 | 122 | 48658 | 00:13:27 | 2158 | 56069 | 00:02:35 | 133 | 66861 |
| 15920441 | 00:00:08 | 237 | 72035 | 00:29:32 | 4096 | 90182 | 00:04:48 | 195 | 132408 |
| 30786116 | 00:00:14 | 454 | 96595 | 01:00:27 | 7575 | 134112 | 00:10:00 | 316 | 256041 |
| 62502135 | 00:00:27 | 917 | 123541 | 02:26:56 | 14234 | 181825 | 00:18:44 | 572 | 519838 |
| 125063199 | 00:00:52 | 1831 | 157818 | 08:43:14 | 24269 | 240747 | 00:38:48 | 1076 | 1040174 |
| 254576853 | 00:01:45 | 3723 | 188813 | >24h | NA | NA | 01:20:03 | 2071 | 2117436 |
| 505422833 | 00:03:31 | 7387 | 223931 | NA | NA | NA | 02:39:03 | 4056 | 4203752 |
| 1003934029 | 00:07:11 | 14673 | 267890 | NA | NA | NA | 05:18:28 | 7980 | 8349888 |
| 2018459352 | 00:15:50 | 29496 | 324797 | NA | NA | NA | 10:30:37 | 15697 | 16788084 |
| 3040260476 | 00:24:52 | 44425 | 366761 | NA | NA | NA | 16:13:43 | 23615 | 25286576 |

**Figure 7:**
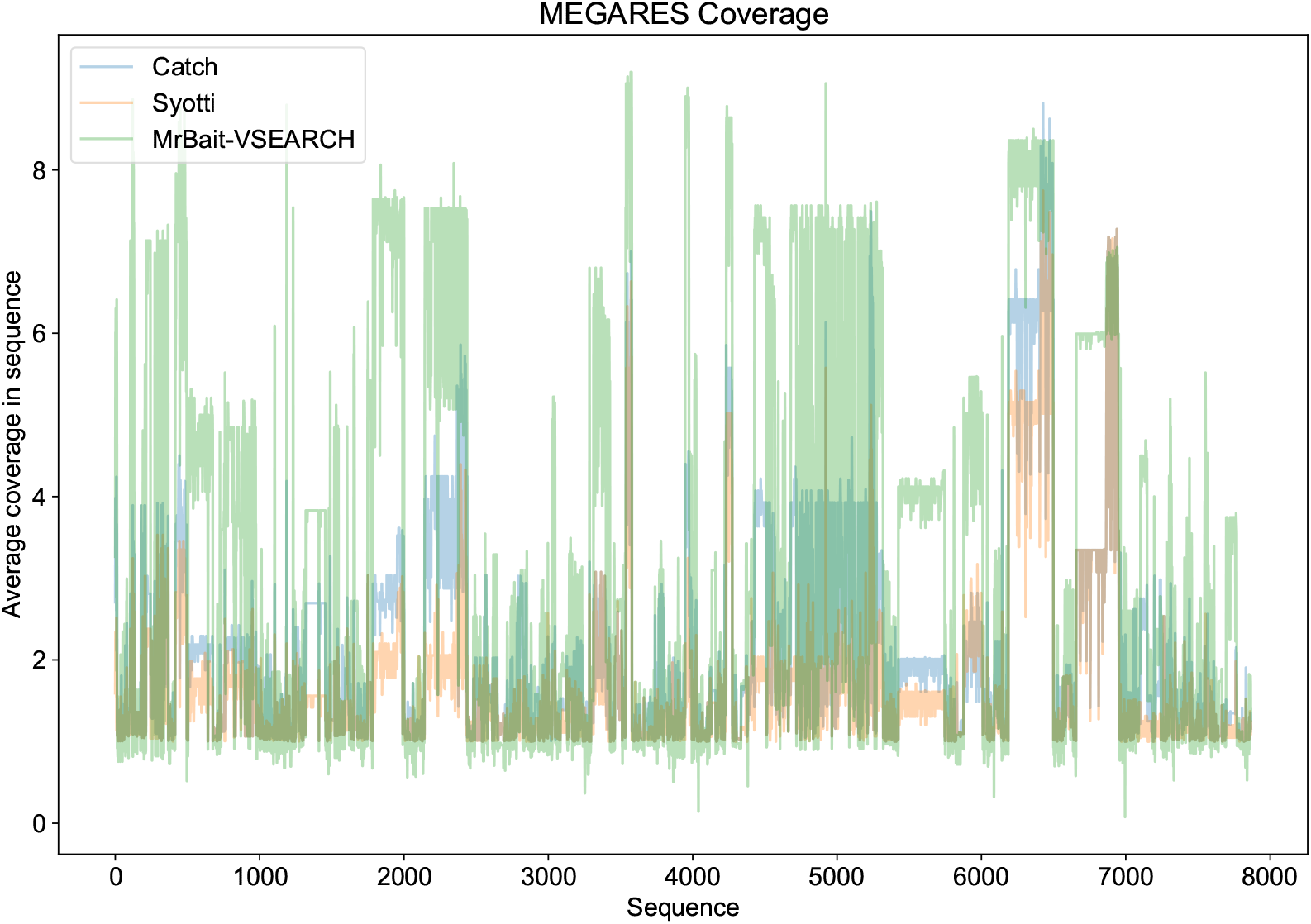
`MEGARES` coverage with VSEARCH filtering for MrBait.

## Footnotes

* Finnish word for “Bait”, with the letter ö replaced with the letter o.

